# Frizzled 6 Drives EMT and Stromal Remodeling Downstream of SP1 in Basal-like Pancreatic Cancer

**DOI:** 10.64898/2025.12.31.697213

**Authors:** Payton D. Stevens, Catriona S. Giannotta, Evan C. Spires, Cassandra R. Diegel, Zhendong A. Zhong, Zachary J. Madaj, Samuel I. Olson, Mohamed H. Dawood, Aaron Honabarger, Giuseppina Caligiuri, David A. Tuveson, Galen B. Hostetter, Bart O. Williams

## Abstract

Pancreatic ductal adenocarcinoma (PDAC) is a highly lethal malignancy, in part due to its rapid progression and early metastasis. To uncover novel drivers of this aggressive behavior, we first analyzed Frizzled receptor expression in human PDAC datasets and identified Frizzled 6 (FZD6) as uniquely predictive of poor clinical outcomes among all ten Frizzled family members. High FZD6 expression correlated with significantly worse overall survival for PDAC patients of any age. Functional studies in human PDAC cell lines showed that FZD6 promotes mesenchymal phenotypes, including enhanced migration and growth in 3D culture, while FZD6 loss induces E-cadherin expression and impairs motility without affecting 2D proliferation. *In vivo*, conditional epithelial deletion of Fzd6 in KPC mice significantly extended survival, reduced tumor burden and metastasis, and remodeled the tumor microenvironment to suppress stromal activation. Single-cell RNA sequencing analysis of human and mouse PDAC tumors confirms that FZD6 is primarily found within the tumor cells and FZD6-activating, noncanonical Wnt ligands are found to be produced by the cancer-associated fibroblasts surrounding the tumors. Mechanistically, we found that FZD6 is a transcriptional target of specificity protein 1 (SP1), a factor upregulated in mutant p53 contexts and within basal-like PDAC subtypes. SP1 knockdown reduced FZD6 expression and signaling through the PCP pathway. Furthermore, reduced SP1 impaired cell motility. Together this implicates the SP1–Fzd6–Wnt/PCP axis as a key driver of epithelial plasticity and disease progression. These findings suggest that FZD6 is a central effector of mutant p53-driven EMT, and a promising therapeutic target in metastatic PDAC.

## Introduction

Pancreatic ductal adenocarcinoma (PDAC) remains one of the most lethal malignancies, with a five-year survival rate of just 13% and limited therapeutic options for most patients at diagnosis (1). A major contributor to PDAC’s poor prognosis is its aggressive biological behavior, particularly the high rate of metastatic dissemination that often accompanies initial clinical presentation. This rapid progression is driven in part by the epithelial-to-mesenchymal transition (EMT), a cellular program that enhances tumor invasiveness, plasticity, and resistance to therapy (2).

The transcription factor specificity protein 1 (SP1) is a critical regulator of EMT in PDAC. SP1 promotes EMT and supports the acquisition of metastatic traits in cancer cells, whereas its inhibition impairs oncogenic signaling and suppresses disease progression (3–5). SP1 expression is notably elevated in genetically engineered mouse models (GEMMs) of PDAC harboring the R172H gain-of-function mutation in Trp53 - a mutation commonly found in human tumors and known to enhance metastatic potential (6,7). Among transcriptional regulators upregulated in response to mutant p53, SP1 is among the most prominent, underscoring its functional importance. Given that TP53 mutations are present in over 75% of PDACs and are enriched in the most aggressive basal-like/squamous subtypes, identifying downstream effectors of SP1 that mediate EMT is essential for developing targeted interventions (8– 12).

Among candidate effectors, recent attention has turned to the Wnt signaling pathway. Mutations in Wnt pathway components are rare in PDAC as they are only present in ∼5% of cases. These mutations primarily affect E3 ligases such as RNF43 and ZNRF3, which ultimately regulate Wnt receptor stability and turnover rates (13,14). Mutations in the E3 ligases increases surface expression of Frizzled receptors, thereby sensitizing cells to Wnt ligands without directly altering canonical or non-canonical signaling proteins (13–16). However, while other components of the Wnt pathway are not commonly mutated, their expression within tumors can vary. Specifically within the non-canonical pathways, others have found that several genes encoding components of the Wnt–Planar Cell Polarity (PCP) pathway are upregulated in metastatic tumors (17).

The PCP pathway, best known for its roles in epithelial sheet organization and oriented tissue morphogenesis, is increasingly implicated in cancer invasion and metastasis. Studies in breast and other epithelial cancers have linked PCP signaling proteins, including Wnt ligands, core receptors, and scaffolding proteins such as Vangl and Prickle, to enhance cell motility and tumor dissemination (18– 22). Frequently, Vangl phosphorylation is a readout for mammalian PCP signaling, which can be activated by several ligands—particularly Wnt4 and Wnt5a—through FZD3 and FZD6 (17,23–27). While FZD3/6 will demarcate the leading edge of migrating PCP cells, cytoplasmic Vangl is phosphorylated and subsequently translocated to the trailing edge of the cells (19,28). Importantly, therapeutic efforts have generally focused on Frizzled receptors, including Vantictumab, a monoclonal antibody that entered clinical trials for PDAC. However, Vantictumab does not bind to FZD6 strongly, which limits its effectiveness for tumors that rely on FZD6-specific signaling (29).

In this study, we identify a transcriptional program driven by SP1 that includes key components of the Wnt–PCP pathway, including FZD6. We show that FZD6 is upregulated in aggressive PDAC subtypes and functionally contributes to EMT and tumor progression. Loss of FZD6 increases epithelial marker expression and impairs cell motility *in vitro. In vivo* loss of Fzd6 delays tumorigenesis in a KPC mouse model, and a comparison of organoid growth in an orthotopic mouse model finds that faster-growing, basal-like organoids have higher levels of PCP receptors. Lastly, analysis of KPC tumors and PDAC patient tumors using scRNAseq highlights that Frizzled 6 is expressed within the tumor cells, and the non-canonical Wnt ligands are expressed within the closely associated cancer-associated fibroblasts (CAFs). These findings position FZD6 as a critical effector of SP1-mediated EMT and a potential therapeutic target in metastatic PDAC.

Importantly, while transcription factors such as SP1 have traditionally been viewed as “undruggable,” targeting cell-surface signaling proteins like Frizzled receptors offers a more accessible route to therapeutic intervention. Importantly, Fzd6-null mice are viable and fertile, exhibiting only hair and nail defects (30,31). In addition, human patients lacking FZD6 also develop nail defects, but otherwise develop normally (32). These data suggest that systemic inhibition of FZD6 may be tolerated with minimal toxicity. This stands in stark contrast to broader Wnt pathway inhibitors, such as Vantictumab and Porcupine inhibitors, which have failed in Phase 1 trials in part due to dose-limiting toxicities, including significant bone pathology (33). Fzd6-null mice do not exhibit bone loss or skeletal abnormalities. Together, this suggests that selective targeting of FZD6 could avoid these adverse effects while still disrupting the pro-tumorigenic EMT program in PDAC.

## Materials and Methods

### Experimental animals

The mice used in this study were maintained in accordance with institutional animal care and use guidelines. Experimental protocols were approved by the Institutional Animal Care and Use Committee (IACUC) of the Van Andel Institute. Mice were fed LabDiet 5021 mouse breeder diet and housed in Thoren Maxi-Miser IVC caging systems with a 12-hour light/12-hour dark cycle. The Frizzled 6 fl/fl, KRAS G12D, Trp53 R172H, and Pdx1-Cre mice were obtained from the Jackson Laboratory (Bar Harbor, Maine). Kras/p53/Pdx1-Cre (KPC) mice on C57BL6 background were maintained by intercrossing to sustain a heterogeneous mixed genetic background. To produce animals used in the experiments, the Frizzled6 fl/fl mice were bred with the KPC mice. These mice were then intercrossed to produce three cohorts of animals, including KPC/Fzd6^+/+^ (Fzd6 WT), KPC/Fzd6^fl/+^, and KPC/Fzd6^fl/fl^ (Fzd6 cKO). To monitor survival, these cohorts of mice were followed for up to 7 months. Mice that presented with either anal or facial papillomas were sacrificed according to the IACUC protocol.

### Cells and reagents

Panc1, BxPC3, MiaPaca-2 cells obtained from American Type Culture Collection (ATCC). Panc1 and MiaPaca-2 cells were cultured in standard DMEM media and supplemented with 10% FBS (Sigma-Aldrich) and 1% penicillin/streptomycin. BxPC3 cells were cultured in RPMI1640 supplemented with 10% FBS and 1% penicillin/streptomycin. Human pancreatic cancer cell lines tested negative for Mycoplasma using PCR (MycoTest Kit).

Stable Frizzled6 knockdown cells were generated by lentivirus-based shRNAs. The shRNA targeting sequences for human Frizzled6 were constructed in pLKO.1-puro vector and purchased from Sigma-Aldrich (34). Two shRNA constructs were used for Frizzled 6 knockdown: TRCN0000008338 (labeled as 8338 in figures) and TRCN0000008341 (labeled as 8341 in figures). A plasmid carrying a scrambled, non-targeting sequence was used to create the control cells. Lentivirus was produced in HEK293T cells by transfecting lentiviral constructs with helper plasmids (pMD2.G (Addgene #12259) and psPAX2 (Addgene # 12260)) using FuGene HD per manufacturer’s instructions. The virus containing media were collected, filtered, and overlaid onto the Panc1, BxPC3, and MiaPaca-2 cells in the presence of polybrene (8 μg/ml) for 24 hours. The infected cells were then subjected to selection with puromycin (4 μg/ml) for one week.

Transient knockdown of SP1 was generated using siRNA dicer-substrate duplexes ordered from Origene. Two constructs targeting SP1 (A and C) were transfected into Panc1, BxPC3, and MiaPaca-2 cell lines. 20nM of each construct was transfected using Origene siTran2.0 reagent into cells grown in serum free OptiMEM. Experiments using these cells occurred within 24 to 48 hours after transient knockdown.

### Transwell migration assay

Transwell migration (or Boyden chamber) assays were performed using standard procedures. Briefly, cell lines were grown to 50% confluency and serum starved overnight. 5 × 10^4^ cells were seeded into Transwell inserts with 8um pores (Corning) in 0.1% BSA and allowed to migrate for 3 - 4 hours at 37°C toward media (DMEM or RPMI1640 depending on the cell line) containing 10% FBS and collagen (15 μg/mL coated on the bottom side of the Transwell insert). At the end of the incubation period, cells migrated to the bottom of Transwell inserts were fixed with methanol for 20 minutes and stained with 0.5% crystal violet. The numbers of cells were counted using an inverted microscope at ×20 magnification.

### Western blot analysis

Pancreatic cancer cells were harvested, lysed (using 50 mM Na2HPO4, 1mM sodium pyrophosphate, 20 mM NaF, 2 mM EDTA, 2 mM EGTA, 1% Triton X-100, 1 mM DTT, 200 µM benzamidine, 40 µg ml^-1^ leupeptin, 200 µM PMSF), and centrifuged (16,000 x g for 3 minutes at 4°C) to obtain detergent-solubilized cell lysates. Equal amounts of cell lysates were resolved by SDS-PAGE and subjected to Western blot analysis, and imaged using the ChemiDoc (BioRad). The Frizzled6 (#5158), SP1 (#9389), E-cadherin (#3195), β-actin (#3700), and GAPDH (#2118) antibodies were from Cell Signaling Technology. The phospho-VANGL2 (p-VANGL2 for Ser79, Ser82, Ser84) was obtained from ThermoFisher (cat#: MA5-38242).

### Orthotopic and RNA sequencing analysis

Orthotopic models and RNA sequencing were performed at Cold Spring Harbor Laboratory (CSHL) where all animal experiments were performed according to procedures approved by the CSHL IACUC (35). Briefly, NOD.Cg-Prkdcscid Il2rgtm1Wjl/SzJ (NSG) mice (Jackson Laboratories) between 3 and 10 weeks of age were used for organoid transplantation. A single-cell suspension of human patient-derived organoids were generated as previously described (36). The intraductal grafting of organoid (IGO) transplantation used 50,000 to 100,000 cells suspended in 25 μl of PBS. These were infused into the pancreatic duct using the modified retrograde infusion technique described previously (37). Following anesthetization with 2.5–3% (weight/volume) isoflurane inhalation, the tumor cells were passed through the small hole that has been created by a needle in the duodenal wall and then into the sphincter of Oddi. A microclamp was placed at the sphincter of Oddi to prevent backflow or leakage during the infusion. Cells were injected by manually depressing the plunger into the syringe for 2-3 minutes. The incision point in the duodenum was closed by suturing (6-0 silk, CP Medical Sutures) or using VETCLOSE surgical glue (Henry Schein CA1999). The pancreas and other organs were carefully placed back into the abdomen before suturing the peritoneal cavity and skin incision (separately).

After PDAC tumors formed the murine pancreas was isolated, minced, and digested for 20 minutes at 37°C in digestion buffer (DMEM, 10% FBS, 1% penicillin, 1% streptomycin, 2.5 mg/ml Collagenase D (Sigma-Aldrich), 0.5 mg/ml Liberase DL (Sigma-Aldrich), 0.2 mg/ml DNase I (Sigma-Aldrich)) with triturating every 5 minutes. After washing with flow buffer (PBS, 2% FBS), cell suspensions were collected and run through 100 μm cell strainers. Red red blood cells were lysed using ACK lysis buffer (Gibco). Cancer cells were enriched with human EpCam-coated beads (130-061-101, Miltenyi Biotec) according to the manufacturer’s instruction. Samples were collected in TRIzol reagent for RNA extraction.

For RNA-seq comparison of Fast-vs. Slow-IGO tumors, 6 IGO transplants of Fast-progressors (hM1A, hF3, hT3, hF24, hF43 and hM19B) and 9 IGO transplants of Slow-progressors (hF2, hF23, hF27, hT30, hF32, hT85, hT93, hT102, and hM17D) were performed, and transplant-derived tumors were harvested at endpoint. RNA from bulk tumors was extracted using the PureLink RNA Mini Kit with TRIzol Reagent as described above.

RNA-seq libraries were constructed using a TruSeq Stranded Total RNA Kit with RiboZero Human/Mouse/Rat (RS-122-2202, Illumina) with 0.2–1 μg RNA per sample, per manufacture’s instructions. Libraries were evaluated using BioAnalyzer (Agilent 2100) and then sequenced using Illumina NextSeq platform (Cold Spring Harbor Laboratory Genome Center, Woodbury).

## Results

### Frizzled 6 Expression Correlates with Poor Prognosis in Human PDAC

To identify Wnt signaling components associated with aggressive pancreatic cancer, we analyzed data from The Cancer Genome Atlas (TCGA) and multiple public expression datasets for human PDAC tumors. Among the ten Frizzled (FZD) family members, FZD6 was most consistently associated with poor clinical outcomes. High FZD6 expression predicted significantly shorter overall survival (OS) in patients with PDAC (**Fig. 1A**). Analysis of CPTAC datasets for top and bottom quartiles of FZD6 expressing PDAC patients further shows the significant impact FZD6 has on patient outcomes, with patients having worse outcomes with higher expression of FZD6. Kaplan-Meier curves for High (red), Mid (orange), or Low (blue) FZD6 are shown in **Fig. 1B**. Further stratifying the patients, we used the combined data from TCGA and CPTAC to analyze the survival probability of patients diagnosed at different ages. While holding age constant, for each transcript per million (TPM) unit increase in FZD6 expression, the hazard ratio increases by ∼127% (95% CI: 12.9% - 357%, p = 0.02) (**Fig. 1C**). In particular, PDAC tumors classified within the squamous or basal-like subtypes—known for aggressive clinical behavior—exhibited the higher FZD6 expression levels. Analysis of Nicolle *et al*. xenograft data (**Fig. 1D**) shows that tumors expressing GATA6, a marker for the less metastatic classical tumors, have lower levels of several PCP proteins (38). Interestingly, after xenograft tumor implantation, the mouse stroma surrounding the more aggressive, basal-like tumors begins to produce more non-canonical stimulating Wnt5a. These data nominate FZD6 as both a prognostic biomarker and a putative functional driver of tumor progression in PDAC.

**Figure 1.**
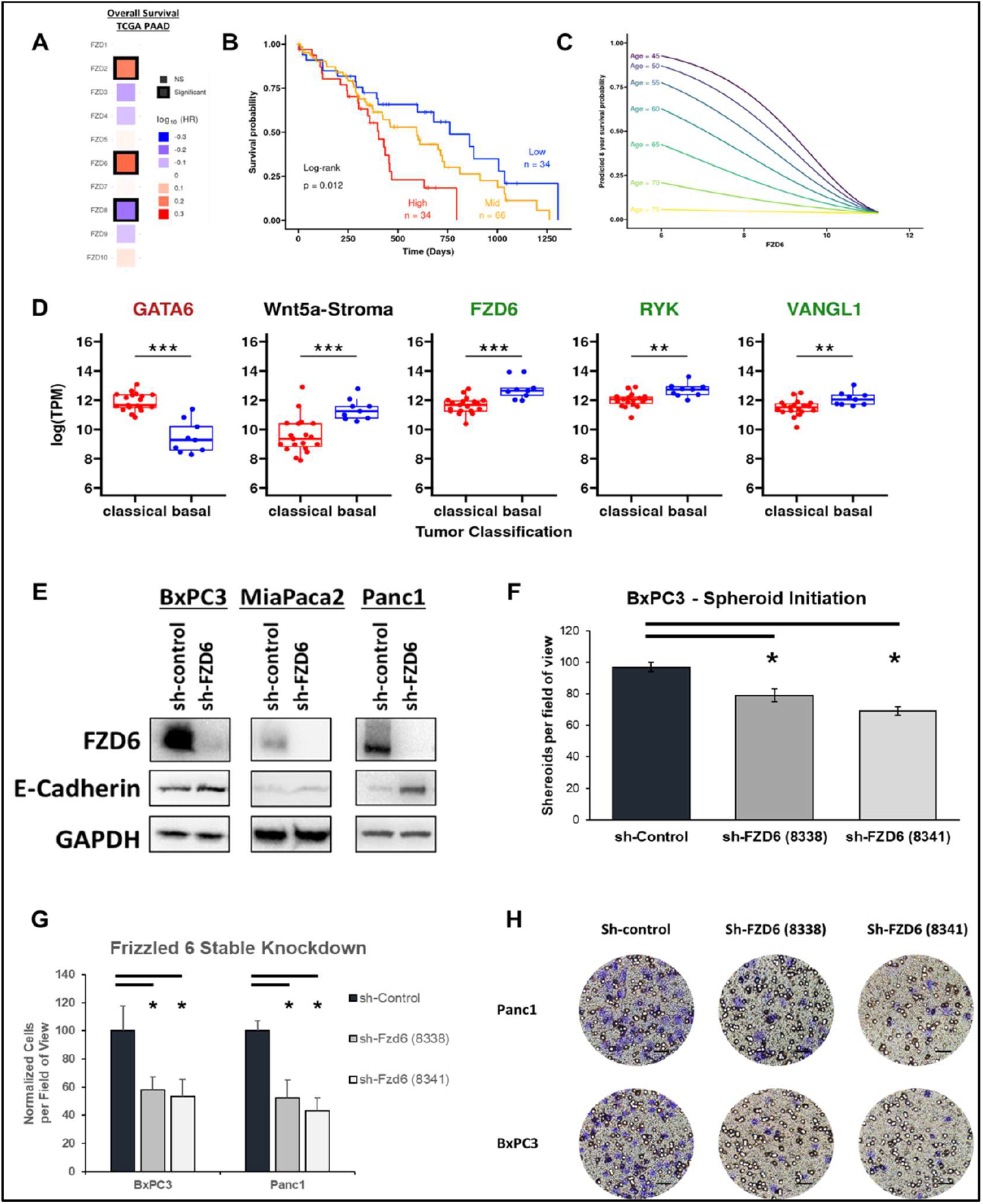
PCP signaling genes are expressed more highly in aggressive PDAC and reduce patient overall survival. **A**. Comparing the expression-to-patient outcome analysis of all ten Frizzled receptors in TCGA PAAD data set. **B**. CPTAC PDAC dataset. Kaplan-Meier survival curves of patients with the lowest 25% of FZD6 expression levels (blue), those between 25% and 75% (yellow), and those with the highest 25% of FZD6 expression levels (red). **C**. Survival probability of patients stratified by age and expression (log2TPM) of FZD6 (TCGA and CPTAC datasets combined). For each unit increase in FZD6 expression, the hazard increased by ∼127% (95% CI: 12.9% - 357%, p = 0.02), holding age constant. **D**. Analysis conducted on subcutaneous Patient-derived xenografts (PDX) dataset from classical or basal-like PDAC samples (#E-MTAB-5039). GATA6 is a biomarker for the classical subtype of PDAC. Wnt5a (in mouse stroma) and FZD6, RYK and VANGL1 in the human cancer cells are all higher in basal-like tumors. **E**. Pancreatic cancer cell lines with stable, sh-RNA knocked-down FZD6 express higher levels of E-cadherin. **F**. Spheroid/colony formation assay - Single cells were seeded in 33% Matrigel® and allowed to grow for 72 hours. Data represents mean +/-SD. *p<0.05 **G**. Transwell migration assay with BxPC3 and Panc1 control and two stable FZD6 knock-down cell lines. Data is presented as a percentage of the migrated control cells. Data represents mean +/-SD. *p<0.05 **H**. Migrated cells were Crystal Violet stained, and cells within four non-overlapping 20X fields of view were counted.

### FZD6 and SP1 Drive EMT and Cell Motility in PDAC Cell Lines

To investigate how FZD6 contributes to cellular plasticity in pancreatic cancer, we used shRNA and siRNA approaches to knockdown FZD6 and SP1 in three PDAC cell lines: BxPC3, MiaPaca2, and Panc1. E-cadherin expression, an epithelial marker often downregulated during EMT, was markedly upregulated following FZD6 knockdown, indicating a shift toward a more epithelial phenotype (**Fig. 1E**). Spheroid, or 3-dimensional growth, was assayed using the PDAC cell lines with FZD6 knockdown. Spheroid initiation and growth was significantly reduced as the tumor cell lines were grown in 33% Matrigel® (**Fig. 1F**). Notably, 2D proliferation was unaffected by FZD6 knockdown (data not shown), indicating that FZD6 is important in regulating the cell-to-cell contact and interactions of 3D culture; and thus, primarily governs invasive behavior and EMT-associated changes rather than general proliferation. In Transwell/Boyden chamber migration assays, knockdown of FZD6 led to a consistent and significant reduction in cell motility across all lines (**Fig. 1G**). Representative images for Panc1 and BxPC3 migration assays are shown (**Fig. 1H**). These results collectively support a model in which FZD6 drives mesenchymal transition, enhances invasive behavior, and potentially contributes to disease aggressiveness in PDAC.

Analysis within Kim *et al*. highlighted the importance of the SP1 transcription factor in the metastatic capabilities of PDAC (**Fig. 2A - left**) (6). KPC tumors harboring p53 R172H mutation, rather than p53 knockout, were ∼40% more metastatic and SP1 was the most significantly increased transcription factor within the aggressive tumors. Supporting this, previous data in PDAC patients have shown that TP53 mutations occur more frequently in the more metastatic, basal-like subset of tumors, mirroring the mouse data in Kim *et al*.(8,9,11). Further analysis of TCGA PAAD data reveals that SP1 transcription is significantly correlated with that of FZD6 (**Fig. 2A, right**). PDAC patients with higher SP1 mRNA expression also have more FZD6. Using siRNA, we found the SP1 knockdown led to a decrease in FZD6 protein levels and decrease in phosphorylated VANGL2, a read-out for active PCP signaling (**Fig. 2B**). The decrease in SP1 within Panc1 and BxPC3 cell lines also reduced cell migration (**Fig. 2C**), as seen in the representative images (**Fig. 2D**). These results suggest that SP1 is a transcriptional activator upstream of FZD6, linking oncogenic TP53 activity to Wnt/PCP signaling and mesenchymal transformation.

**Figure 2.**
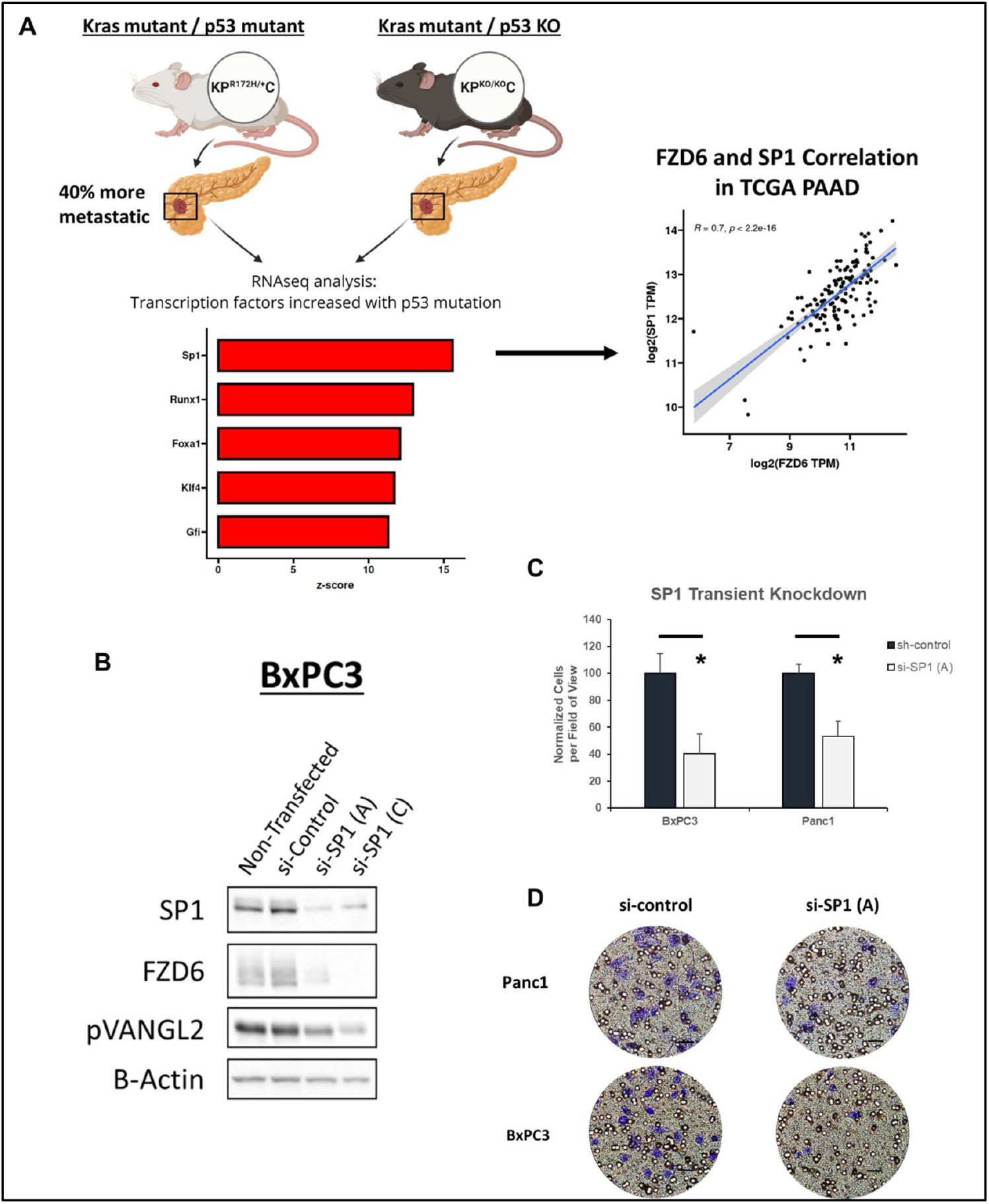
Reduction of SP1 expression reduces PCP signaling and PDAC cell motility. **A**. Data from Kim *et al*. showed SP1 transcription factor is the most highly induced after R172H mutation in KPC mice (compared to p53 null KPC mice); TCGA analysis of SP1 expression is highly correlated with FZD6 **B**. Use of siRNA for transcription factor SP1 results in reduced expression of FZD6 and decreased phosphorylation of VANGL2. **C**. Transwell migration analysis after knockdown of SP1 results in reduced cell motility Data represents mean +/-SD. *p<0.05. **D**. Representative images from Transwell chambers showing a trend in decreased migration for Panc1 and BxPC3 cell lines upon SP1 knockdown.

### Epithelial Fzd6 Deletion Extends Survival and Suppresses Tumor Progression in KPC Mice

To evaluate the *in vivo* role of Fzd6 in PDAC, we generated KPC mice with epithelial-specific deletion of Fzd6 (KPC-Fzd6^cKO^) and compared them to littermate controls (KPC-Fzd6^WT^). Kaplan–Meier survival analysis showed that Fzd6^cKO^ mice survived significantly longer than Fzd6^WT^ counterparts (**Fig. 3A**). Mice that were heterogeneous for the Fzd6 allele, while also significantly longer lived than their KPC-Fzd6^WT^ littermates, were found to be intermediate of the KPC-Fzd6^WT^ and KPC-Fzd6^cKO^. At necropsy, Fzd6-deficient tumors were smaller and less likely to exhibit gross metastasis: 28% (6/21) of KPC-Fzd6^WT^ mice had visible metastatic lesions, compared to only 5% (1/17) of KPC-Fzd6^cKO^ animals (**Fig. 3B**). The single Fzd6^cKO^ animal with metastasis was a 7-month-old mouse, significantly older than the typical endpoint for KPC mice, suggesting delayed disease progression.

**Figure 3.**
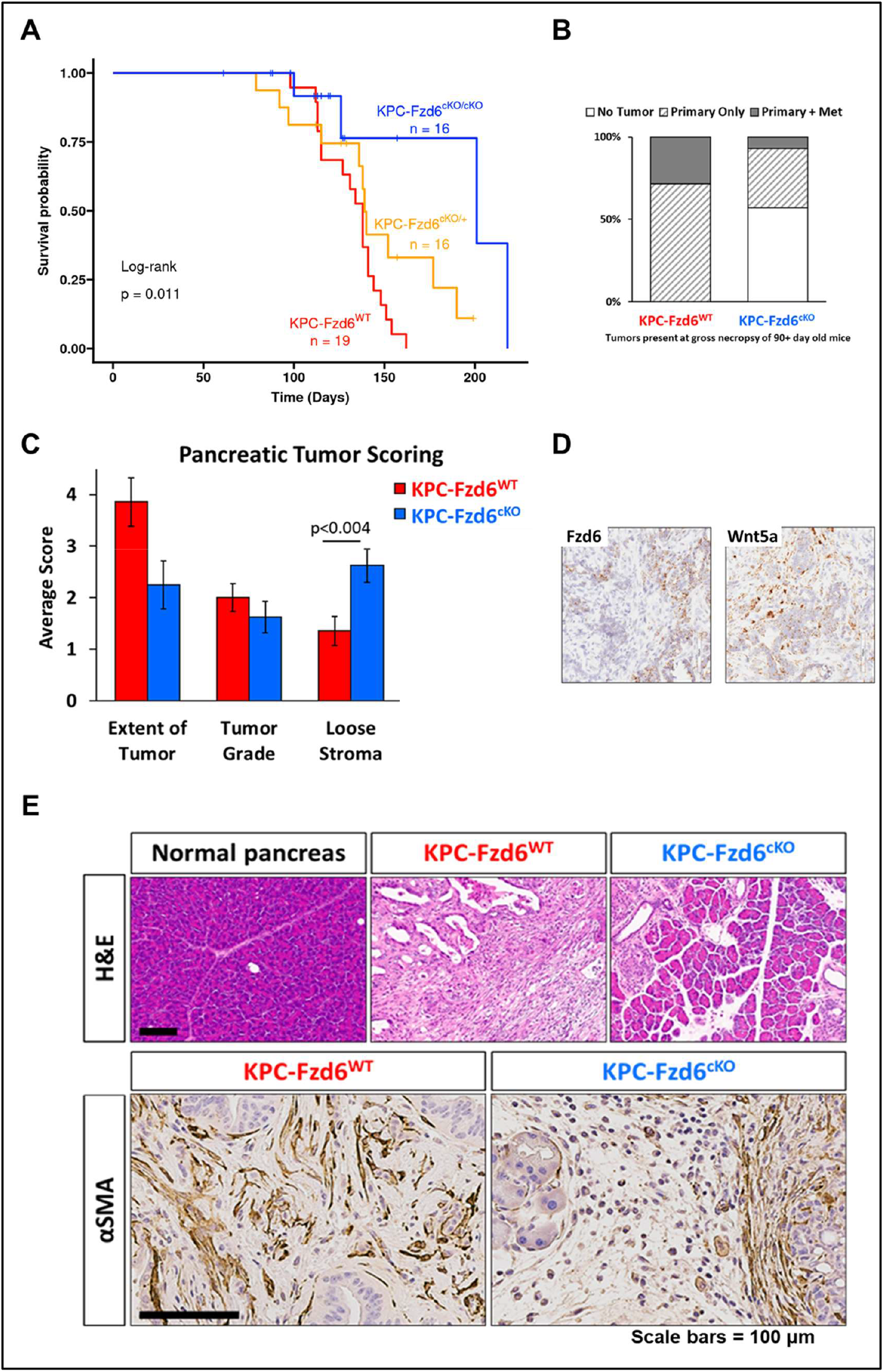
Loss of Fzd6 inhibits PDAC progression and alters the tumor microenvironment. **A**. Kaplan-Meier plot for KPC-Fzd6^WT^, KPC-Fzd6^cKO/+^,and KPC-Fzd6^cKO/cKO^. **B**. Quantification of grossly observed tumors during necropsy of mice older than 90 days of age. Note the only KPC-Fzd6^cKO^ mouse with a metastasis was over 7 months old. **C**. Pathologist provided histological analysis of PDAC in KPC-Fzd6^cKO^ mice (Blinded Scoring) **D**. KPC tumors stained using RNAscope show Fzd6 and Wnt5a expression paterns **E. Top**. H&E-stained sections of pancreas tissue from a wild-type mouse, pancreas tissue overtaken by PDAC in a KPC-Fzd6^WT^ mouse, and a pancreas from a KPC-Fzd6^cKO^ mouse. **Bottom**. αSMA images of PDAC tissue in KPC-Fzd6^WT^ and KPC-Fzd6^cKO^ mice. Scale = 100µm

KPC mice also occasionally develop facial and anal papillomas (requiring the mice to be sacrificed prematurely) due to the off-target activity of Pdx1-Cre. These off-target effects develop at the same rate regardless of Fzd6 status, suggesting that the tumor-inhibiting effects of Fzd6 deficiency are specific to PDAC, at least relative to the cell of origin for the papillomas. Importantly, we found that among KPC animals sacrificed after 90 days of age, Fzd6 deletion consistently reduced tumor burden and dissemination, without increasing the frequency of off-target tumor phenotypes.

### Fzd6 Loss Remodels the Tumor Microenvironment and Restrains Stromal Activation

To understand the impact of epithelial-specific Fzd6 loss on the pancreatic tumor microenvironment (TME), we performed histological and immunohistochemical analyses on pancreatic tissues from KPC-Fzd6^WT^ and KPC-Fzd6^cKO^ mice. Hematoxylin and eosin (H&E) staining of tumor sections revealed notable differences in tissue architecture. To quantify these morphological changes, we employed a standardized histopathological scoring system across three metrics: tumor extent, tumor grade, and stromal architecture (**Fig. 3C**). To confirm localization of Wnt pathway proteins within the KPC tumor and microenvironment RNAscope® was performed (**Fig. 3D**), with Fzd6 demarcating the tumor-epithelial cells and Wnt5a within the cells immediately surrounding the tumors.

As expected from histopathological analysis, the KPC-Fzd6^WT^ tumors exhibited dense, poorly organized neoplastic regions consistent with late-stage PDAC. In contrast, tumors from KPC-Fzd6^cKO^ animals retained a more preserved glandular architecture with reduced parenchymal disruption, suggesting delayed or diminished progression (**Fig. 3E - top**). Notably, KPC-Fzd6^cKO^ tumors exhibited a significant reduction in overall tumor burden and invasiveness compared to their Fzd6^WT^ counterparts. In addition, the stroma in Fzd6-deficient tumors was more loosely organized and less ECM-rich (p < 0.05), both of which are features that have previously been associated with increased therapeutic accessibility.

Further insight into stromal remodeling was gained through α-smooth muscle actin (α-SMA) staining, a marker of activated CAFs (**Fig. 3E - bottom**). In KPC-Fzd6^WT^ tumors, α-SMA expression was widespread and intense, consistent with an abundant and contractile stromal response. However, in KPC-Fzd6^cKO^ tumors, α-SMA staining was substantially reduced and reorganized, indicating a potential shift in CAF activation status or subtype composition in the absence of epithelial Fzd6.

### FZD6 PCP Signaling Is Conserved in Patient-Derived PDAC Organoid Models

We next examined gene expression in organoids derived from the Tuveson laboratory IGO orthotopic model of PDAC (**Fig. 4A - left**) (39). This model allows for human patient-derived PDAC organoids to be transplanted into the pancreatic duct of immunodeficient mice. Importantly, compared to other orthotopic transplant models, this specific placement of the organoid tumor cells allows them to maintain their classical or basal-like characteristics. Therefore, prior to orthotopic implantation the organoids were stratified by epithelial, classical (Slow-IGO) versus mesenchymal, basal-like (Fast-IGO) subtypes. The organoids were transplanted and allowed to grow, followed by RNA sequencing of the resulting tumors. Within the patient organoids, Fzd6 and the PCP co-receptor RYK were enriched in the Fast IGO, basal-like subset (**Fig. 4A, right**). Additionally, similarly to the xenograft data from Nicolle *et al* (**Fig. 1D**), the transplantation of basal-like human tumor cells resulted in a change to the surrounding mouse stroma. Specifically, the mouse stroma increased the production of non-canonical Wnt ligand, Wnt5a. These data support that Fzd6 and PCP signaling are predominant in aggressive PDAC subtypes.

**Figure 4.**
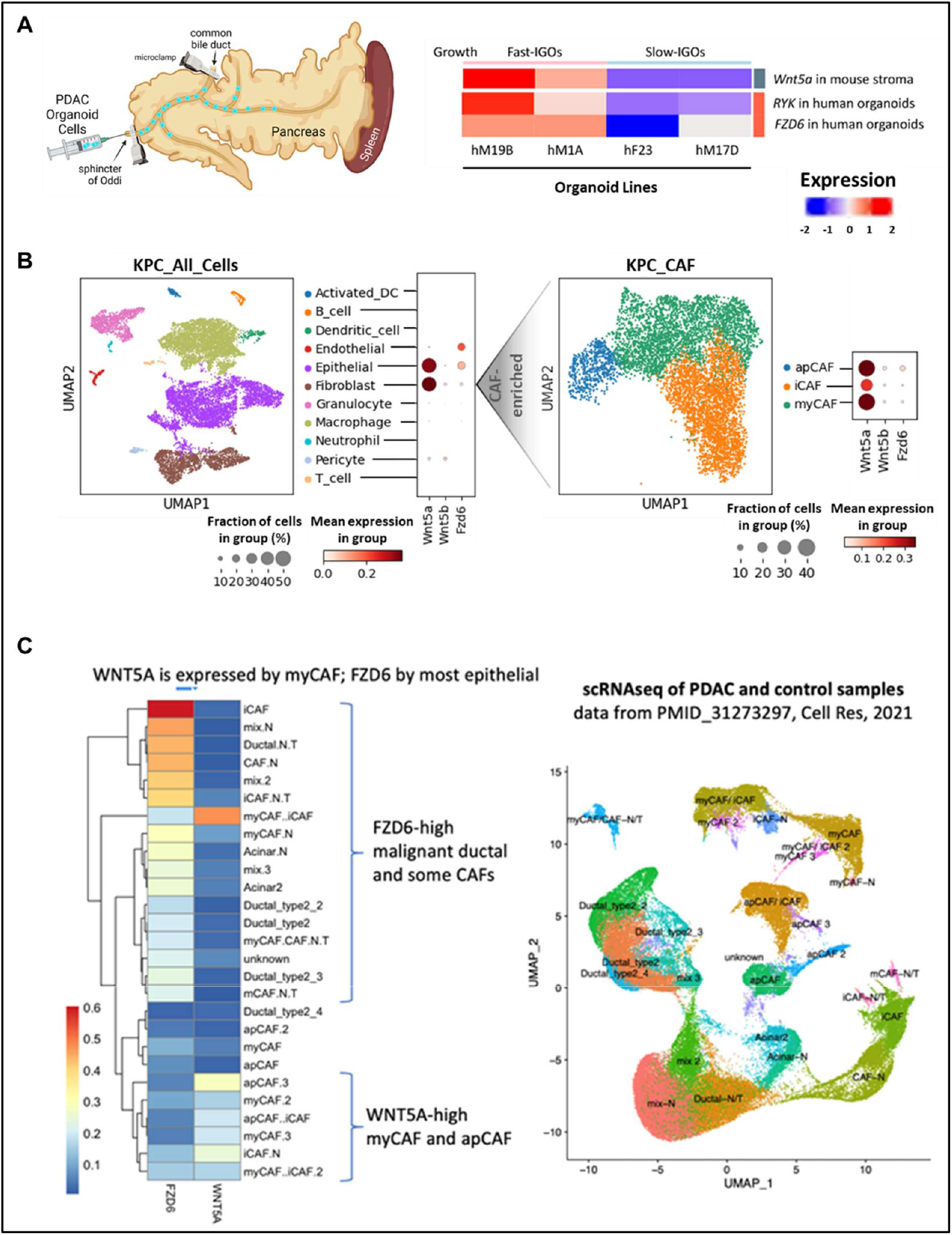
Epithelial FZD6 and fibroblast WNT5A are found in patient tumor organoids. **A. Left**. Model depicting the Intraductal Grafted Organoid (IGO) procedure; the injection of human organoid cell suspension into murine pancreatic ducts. Created with BioRender.com **Right**. The expression of FZD6 and RYK in human PDAC cells and Wnt5a in mouse stroma were assessed by RNA sequencing. FZD6, and co-receptor RYK, are more highly expressed in fast-growing human-organoid derived tumors. Wnt5a is more highly expressed in fast-IGO mouse stroma compared to stroma surrounding slow-IGO tumors. **B. Left**. scRNAseq was performed on all cells from a KPC tumor. **Right**. Fibroblast-enriched tumor cells were sorted DAPI-;CD45-;CD31-; EPCAM- and further analyzed for gene expression within the murine cancer-associated-fibroblasts (CAF) subtypes. **C**. scRNAseq analysis of Peng *et al*. compares patient PDAC tumors to normal pancreas and shows FZD6 and WNT5A localization within the tumor.

To confirm the expression of FZD6 within PDAC tumor cells and the supportive role of the tumor stroma through Wnt ligand secretion, scRNAseq analysis was performed on KPC and patient tumor cells. From KPC tumor analysis we found that Fzd6 was primarily limited to the epithelial and endothelial cells of the tumor (**Fig. 4B - left**). Additionally, as we focused on Wnt ligand secretion, we confirmed that the tumor cells, as well as myofibroblastic CAFs (myCAFs) and antigen-presenting CAFs (apCAFs), secrete the non-canonical Wnt5a ligand. The tumor constraining myCAFs are closest in proximity to the epithelial cells within a KPC tumor, and apCAFs are typically distributed throughout the TME. Confirming the role of PCP signaling in patient data – scRNAseq from Peng *et al*. was analyzed. These data used single-cell transcriptomes of ∼2,000 PDAC cells and ∼15,000 normal cells from control pancreases (40). FZD6 was found to be highly expressed within the epithelial compartment of the patient tumors, with some expression in specific CAF subsets. WNT5A in the patient cells was found to be expressed similarly to Wnt5a in KPC tumors, with the highest expression observed within myCAF and apCAF populations (**Fig. 4C**).

Together, these results demonstrate that epithelial FZD6 not only drives tumor cell-autonomous progression but also profoundly influences the organization and functional state of the TME. The findings suggest that FZD6 regulates epithelial–stromal signaling in a manner that promotes desmoplasia, a hallmark of the aggressive basal-like PDAC subtype. These data strongly support our central hypothesis that the Fzd6–Wnt/PCP signaling orchestrates epithelial plasticity and stromal remodeling in pancreatic cancer.

## Discussion

Our findings establish FZD6 as a critical mediator of EMT, tumor progression, and stromal remodeling in PDAC, acting downstream of SP1 and within a Wnt/PCP signaling context. High FZD6 expression, unique among Frizzleds, predicts poor clinical outcomes across PDAC datasets and is most enriched in the basal-like subtype - known for high-grade tumors, mutant p53 expression, and aggressive clinical behavior. Through *in vitro, in vivo*, and organoid studies, we demonstrate that FZD6 promotes tumor intrinsic, invasive and mesenchymal characteristics while also shaping tumor extrinsic characteristics, such as altering the TME in ways that support stromal restructuring and desmoplasia.

Both SP1 and FZD6 loss led to impaired cellular motility, with FZD6 reduction leading to re-expression of epithelial markers, which are consistent with their roles in EMT. This convergence of phenotypic effects reinforces a hierarchical model in which SP1 transcriptionally activates FZD6 to initiate Wnt/PCP signaling, promoting mesenchymal differentiation (3,17). The coordinated suppression of both transcriptional and signaling components of this axis effectively impairs the acquisition of invasive phenotypes. Additionally, our results align with a broader body of literature that links SP1 activity to basal-like tumor states in PDAC, suggesting that targeting SP1 or its downstream effectors like FZD6 could yield subtype-specific therapeutic vulnerabilities (3,4). Others have noticed that mutant p53 status can result in more metastasis, and alter CAFs and stroma within the TME (6,41). We hypothesize that the mechanism is at least partially driven by the SP1-FZD6 signaling. These data suggest a model in which SP1, activated by oncogenic KRAS and p53 signaling, drives a mesenchymal program via FZD6 induction, with downstream engagement of non-canonical Wnt/PCP signaling cascades.

Our murine genetic data further solidify the *in vivo* relevance of this pathway. Epithelial-specific Fzd6 deletion in KPC mice significantly extended survival, reduced metastatic dissemination, and dramatically altered the TME. In particular, Fzd6-deficient tumors showed reduced fibroblast activation, a hallmark of a more therapy-permissive stroma. These findings provide new insights into how EMT regulators like Fzd6 may influence not only cancer cell-intrinsic behavior but also cell extrinsic behaviors, such as the composition and function of the TME. Interestingly, these data implicate the role of PCP signaling in changing the tumor secretome, which would allow the tumor cells to have such a large, wide-spread impact. Additionally, while tumor-cell intrinsic changes potentially allow FZD6-rich cells to be motile and leave the primary tumor site, we hypothesize the observed tumor extrinsic changes may alter the tumor immune landscape. Several of the TME alterations we observed are known to impact immune cell infiltration and activation, leading to intriguing future research directions.

Given that traditional EMT factors such as transcriptional repressors are difficult to target pharmacologically, FZD6 represents a tractable point of intervention. As a cell-surface receptor with limited roles in adult homeostasis, targeting FZD6 could impair tumor progression and EMT without inducing broad toxicity. The viability of Fzd6-null mice supports this proposition.

Together, our work positions FZD6 as a functionally validated EMT effector, transcriptionally regulated by SP1, and a key orchestrator of tumor–stromal crosstalk in aggressive pancreatic cancer. Therapeutic inhibition of FZD6 could simultaneously blunt EMT, improve the efficacy of existing therapies, and/or potentially sensitize tumors to immune attack. Notably, earlier attempts to therapeutically target Frizzled receptors, such as the use of Vantictumab in clinical trials, were not effective in PDAC, perhaps because of the low affinity of Vantictumab for FZD6 (33). Our findings suggest that a more targeted strategy focusing specifically on FZD6 may yield greater efficacy in tumors where this receptor is the principal effector of Wnt/PCP-driven EMT and metastasis.

## Acknowledgements

We thank members of the Williams and Tuveson labs for helpful discussions. We are grateful to the VAI Vivarium and Histology Core for expert technical assistance, and to the Cold Spring Harbor Laboratory for sharing organoid models. We also thank patients and families who contributed to public datasets used in this study.

This work utilized the following Van Andel Institute core facilities, supported in part by institutional funding and the respective RRIDs: Pathology and Biorepository Core (RRID:SCR_022937), Optical Imaging Core (RRID:SCR_022939), Bioinformatics and Biostatistics Core (RRID:SCR_022940), Vivarium Core (RRID:SCR_022938), and Transgenic Core (RRID:SCR_022941).

Images were created using Biorender.com. Grammarly and ChatGPT were used for editing.

